# Fraction of copy-number alterations significantly predicts survival following immunotherapy in a few cancers

**DOI:** 10.1101/2022.12.28.522101

**Authors:** Tiangen Chang, Yingying Cao, Eldad D. Shulman, Alejandro A. Schäffer, Eytan Ruppin

## Abstract

Various studies have shown that high tumor mutation burden (TMB) may predict response to immune checkpoint therapy, at least in some cancer types ^1,2^. However, identifying patients with low TMB that are still likely to respond to cancer immunotherapy is an important open challenge. Recently, Spurr et al. ^3^ reported that the tumor *aneuploidy score* (AS), defined as the fraction of chromosome arms with arm-level copy number alterations in a sample, is predictive of survival following immunotherapy in low-TMB patients across multiple cancers. By re-analyzing the same data set by performing survival analysis in individual cancer types separately, we find that AS only significantly predicts survival in one single cancer indication. We further find that another metric conceptually related to the AS, *the fraction of genome encompassed by copy number alterations (FGA)*, if called with a conventional copy number calling cutoff, has stronger predictive power than the AS proposed in ^3^, and that this observation holds even if the FGA and AS thresholds for presence/absence of copy number events are set comparably. However, with the current available data, even FGA can predict survival following immunotherapy in only a few cancer indications.

---

One promising approach to identify low-TMB responders of immunotherapy has been to study the predictive ability of other measures of genomic alterations in cancer in these patients. Two natural candidates are (a) tumor aneuploidy, which measures chromosome-level copy number alterations, and (b) genomic copy number alterations, which quantifies the extents of both chromosomal and focal copy-number events ^4^. Both tumor aneuploidy and genomic copy number alterations have been shown to play a role in cancer progression and to be predictive to cancer prognosis ^4-6^. Recently, Spurr et al. ^3^ analyzed data of an MSK-IMPACT cohort (Samstein et al. cohort) ^1^, reporting data and response information about 1,660 advanced cancer patients from 10 different cancer types treated with immune checkpoint blockade (ICB). They reported that in low-TMB patients (the bottom 80% within each cancer type), AS significantly predicted *pan-cancer* overall survival following immunotherapy, by aggregating the data across all cancer types. Intrigued by their potentially clinically impactful finding, we set out to explore two related fundamental questions in the same MSK-IMPACT data: (1) Is AS predictive of ICB response of low-TMB patients following immunotherapy in *individual* cancer types? (2) Is AS really a better predictor than FGA?

Firstly, as a pan-cancer Kaplan-Meier survival analysis (what Spurr et al. did in ^3^) may be confounded by factors such as differences in patient numbers in cancer-type composition of the overall dataset when studied in aggregate, we compared the survival curves of low-TMB patients with high versus low AS for each of the 10 cancer types individually. The initial cancer-type-specific analysis was done by using the AS values provided by ^3^, calling chromosome-level copy number alterations by using a cutoff of |log2 copy ratio| > 0.1 (denoted as AS0.1). As evident, in low-TMB patients having low AS0.1, there is a statistically significantly better survival after immunotherapy in only one individual cancer type, “cancer of unknown primary” (n = 70, HR = 2.27, p = 0.031; **Fig. 1**). Here, the hazard ratio (HR) denotes the relative risk of the high-AS0.1 individuals compared to the low-AS0.1 set as the reference.

**Figure 1.**
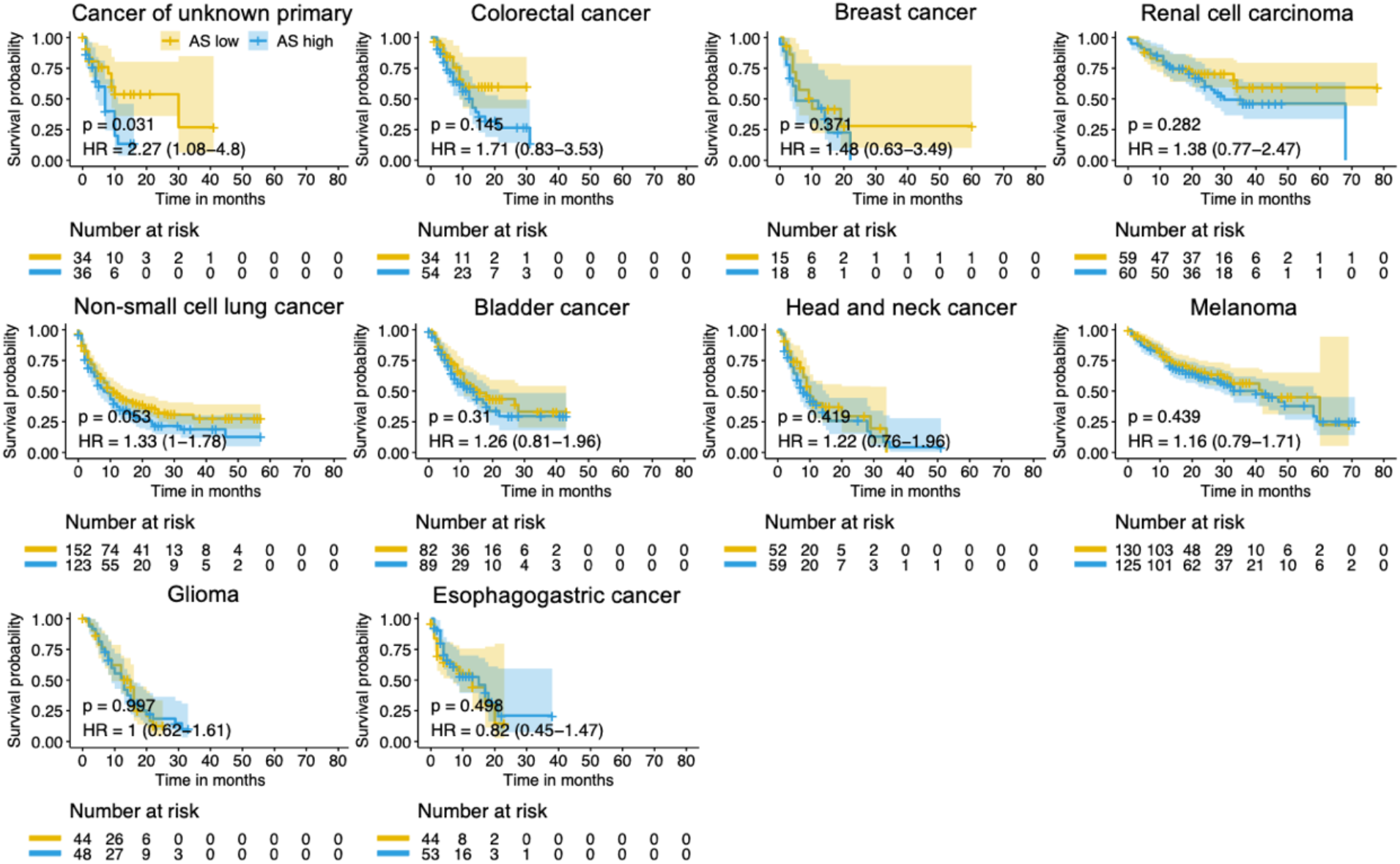
Tumor aneuploidy score (AS0.1) significantly predicts survival following immunotherapy in only one single cancer indication (cancer of unknown primary). Kaplan-Meier survival curves following immunotherapy are compared for low-TMB patients with high versus low AS0.1 (binned at the 50th percentile) in all 10 individual cancer types. AS0.1 is calculated while maintaining the same cutoff as in ^3^ (|log2 copy ratio| > 0.1). Two-sided log-rank p values are indicated, with univariate Cox regression HR with 95% confidence interval. The HR denotes the relative risk of the high-AS0.1 individuals compared to the low-AS0.1 set as the reference.

Next, given that FGA combines both chromosomal and focal copy number alterations and is a higher-resolution measure than tumor aneuploidy, we turned to re-examine the original conclusion of ^3^ that AS is a better predictor than FGA in low-TMB patients. Their analysis was based on using a single cutoff to determine copy number alteration events, |log2 copy ratio| > 0.1 (denoted as FGA0.1). However, other studies, such as ^7,8^, had commonly used the cutoff of |log2 copy ratio| > 0.2. We hypothesized the low cutoff used in ^3^ might be suboptimal, thus dampening the predictive power of FGA and possibly also that of AS. We hence repeated the analysis of ^3^ with the additional conventionally used cutoff of 0.2 (denoted as FGA0.2). Firstly, FGA0.2 has a considerably stronger predictive power for pan-cancer overall survival in low-TMB patients after immunotherapy (n = 1,311, HR = 1.35, p-value = 0.00008) than the original AS0.1 (HR = 1.25, p-value = 0.003) (**Fig. 2a, b**). Secondly, employing a univariate Kaplan-Meier survival analysis in low-TMB patients, we find FGA0.2 significantly or marginally significantly predicts survival in cancer of unknown primary (n = 70, HR = 2.79, p = 0.007), renal cell carcinoma (n = 119, HR = 2.2, p = 0.012), melanoma (n = 255, HR = 1.57, p = 0.023), bladder cancer (n = 171, HR = 1.57, p = 0.048), and non-small cell lung cancer (n = 275, HR = 1.3, p = 0.073) (**Supplementary Fig. 1**). Thirdly, we performed a multivariate overall survival analysis in individual cancer types using Cox proportional-hazards regression. FGA0.2 is significantly or marginally significantly associated with overall survival in renal cell carcinoma (p = 0.02), melanoma (p = 0.003), bladder cancer (p = 0.012), and non-small cell lung cancer (p = 0.077), with adjustment for TMB and drug class of ICB (**Fig. 2c**). This result is consistent with the cancer-type-specific univariate analysis mentioned above. In contrast, none of the multivariate hazard ratios were significant in individual cancer types for AS0.1 measure proposed in ^3^ (**Fig. 2d**). For comparison, we also tested the performance of AS calculated with cutoff of |log2 copy ratio| > 0.2 (denoted as AS0.2). AS0.2 achieves better performance than AS0.1, testifying to the overall better power of this event calling threshold, but it is still less predictive than FGA0.2 (**Supplementary Fig. 2**).

**Figure 2.**
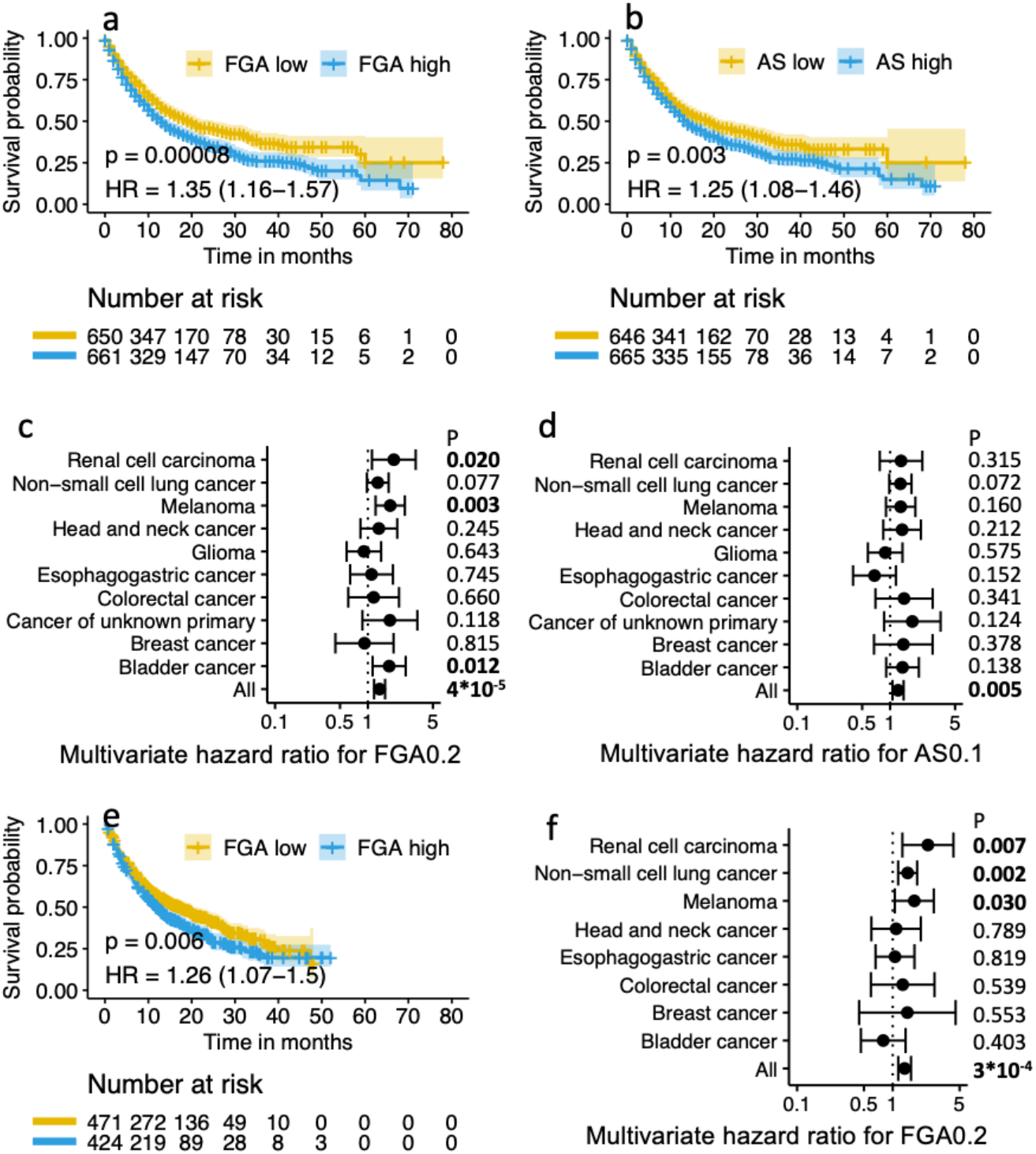
The fraction of copy number alterations (FGA0.2) outperforms tumor aneuploidy score proposed in ^3^ (AS0.1) in predicting survival following immunotherapy. FGA0.2 is re-calculated with a cutoff of |log2 copy ratio| > 0.2. AS0.1 is calculated while maintaining the same cutoff as in ^3^ (|log2 copy ratio| > 0.1). The performance of FGA0.2 and AS0.1 is compared based on both a pan-cancer univariate Kaplan-Meier survival analysis **(a-b)** and via a cancer-type-specific multivariate Cox proportional-hazards regression **(c-d)**. The performance of FGA0.2 is further tested and validated on the Chowell et al. cohort ^8^ (**e-f**). In panels **a, b, e**, two-sided log-rank p values are indicated, with univariate Cox proportional-hazards regression HRs with 95% confidence intervals. In panels **c, d, f**, multivariate HRs are calculated with Cox proportional hazards regression of overall survival using FGA, TMB, and drug class (FGA0.2 binned at the 50th percentile and TMB binned at the 20th percentile), and Wald test multivariate adjusted p values are displayed. The univariate and multivariate survival analysis are done following ^3^ as is.

To further test the robustness of FGA0.2 in predicting survival after immunotherapy in other datasets, we re-analyzed another MSK-IMPACT cohort, which was published recently by Chowell et al. ^8^. In the Chowell et al. cohort, there are eight cancer types in common with the Samstein et al. cohort. We find that the FGA0.2 significantly predicts pan-cancer overall survival in low-TMB patients in the univariate Kaplan-Meier survival pan-cancer analysis (n = 895, HR = 1.26, p-value = 0.006; **Fig. 2e**). When tested in individual cancer types, FGA0.2 significantly or marginally significantly predicts overall survival for three cancer types in this cohort, including renal cell carcinoma (n = 68, HR = 2.44, p-value = 0.012), melanoma (n = 145, HR = 1.87, p-value = 0.018), and non-small cell lung cancer (n = 427, HR = 1.25, p-value = 0.065), but does not significantly predict survival in other cancer types (**Supplementary Fig. 3**). This result is aligned and further confirms the results found in the Samstein et al. cohort (except for bladder cancer; **Supplementary Fig. 1**). Performing a multivariate overall survival analysis in individual cancer types using Cox proportional-hazards regression in this independent dataset, we find that FGA0.2 is significantly associated with overall survival in renal cell carcinoma (p = 0.007), non-small cell lung cancer (p = 0.002), and melanoma (p = 0.03), with adjustment for TMB and drug class of ICB (**Fig. 2f**). This result is again consistent with the cancer-type-specific univariate analysis mentioned above.

In summary, we have comparatively assessed the power of AS and FGA in predicting low-TMB patient survival after immunotherapy in individual cancer types. Addressing our research questions, we first show that the original AS presented in ^3^ (AS0.1) predicts significant survival benefit after immunotherapy in low-TMB patients in only one single cancer type when assessed via a univariate Kaplan–Meier analysis and in none of the cancer types when assessed via a multivariate analysis. Second, in difference from the findings of ^3^, FGA0.2, re-calculated using a conventionally used and more conservative calling cutoff, has a considerably stronger predictive power than AS0.1. Importantly from a translational standpoint, the currently available data suggest that even FGA0.2 can only significantly predict survival following immunotherapy in certain cancer types. The pan-cancer findings of ^3^ are an important and encouraging contribution, but much more data needs to be collected and analyzed before the use of these measures in specific cancer types can be further ascertained, as one hopes.

## Methods

Data for the Samstein et al. ^1^ cohort (MSK-IMPACT) were downloaded and processed following the method described in ^3^ exactly.

Kaplan–Meier survival analysis was performed using the R package “survfit” ^9^, and log-rank P values are reported. Multivariate analysis was performed with Cox proportional hazard regression in individual cancer types using the R package “coxph” ^9^, with inclusion of variates including FGA, TMB and ICB drug class (FGA binned at the 50th percentile and TMB binned at the 20th percentile).

## Data availability

Data for the Samstein et al. cohort are available at https://www.cbioportal.org/study/summary?id=tmb_mskcc_2018 and the GENIE ^10^ v.7.1 release: https://www.synapse.org/#!Synapse:syn7222066/wiki/405659. Data for the Chowell et al. cohort are available from the supplementary table of ^8^, where FGA, TMB, drug class of ICB, and overall survival information are provided. Aneuploidy scores were called using ASCETS at https://github.com/beroukhim-lab/ascets and values for each sample are provided in the GitHub repository referenced below. Source data are provided with this paper.

## Code availability

All code necessary to replicate these analyses are provided in the following GitHub repository: https://github.com/rootchang/Aneuploidy-FGA-ICB.

## Author contributions

T.C. and E.R. conceived and designed the study. T.C. and Y.C. collected and managed the data. T.C., Y.C. and E.D.S. performed the statistical analyses. A.A.S. provided statistical advice. All authors critically revised the manuscript for important intellectual content.

## Acknowledgments

This research was supported in part by the NIH Intramural Research Program, National Cancer Institute. This work utilized the computational resources of the NIH HPC Biowulf cluster (http://hpc.nih.gov). The authors would like to acknowledge the American Association for Cancer Research and its financial and material support in the development of the AACR Project GENIE registry, as well as members of the consortium for their commitment to data sharing. Interpretations are the responsibility of study authors.

## References

1. Samstein, R.M. et al. Tumor mutational load predicts survival after immunotherapy across multiple cancer types. Nature Genetics 51, 202-+ (2019).

2. McGrail, D.J. et al. High tumor mutation burden fails to predict immune checkpoint blockade response across all cancer types. Annals of Oncology 32, 661–672 (2021).

3. Spurr, L.F., Weichselbaum, R.R. & Pitroda, S.P. Tumor aneuploidy predicts survival following immunotherapy across multiple cancers. Nat Genet 54, 1782–1785 (2022).

4. Ben-David, U. & Amon, A. Context is everything: aneuploidy in cancer. Nature Reviews Genetics 21, 44–62 (2020).

5. Hieronymus, H. et al. Tumor copy number alteration burden is a pan-cancer prognostic factor associated with recurrence and death. Elife 7(2018).

6. Sansregret, L. & Swanton, C. The Role of Aneuploidy in Cancer Evolution. Cold Spring Harbor Perspectives in Medicine 7(2017).

7. Spurr, L.F. et al. Quantification of aneuploidy in targeted sequencing data using ASCETS. Bioinformatics 37, 2461–2463 (2021).

8. Chowell, D. et al. Improved prediction of immune checkpoint blockade efficacy across multiple cancer types. Nature Biotechnology 40, 499-+ (2022).

9. Therneau, T. A package for survival analysis in S. R package version 2(2015).

10. Andre, F. et al. AACR Project GENIE: Powering Precision Medicine through an International Consortium. Cancer Discovery 7, 818–831 (2017).

