## Supplementary information for "Fraction of copy-number alterations significantly predicts survival following immunotherapy in a few cancers"

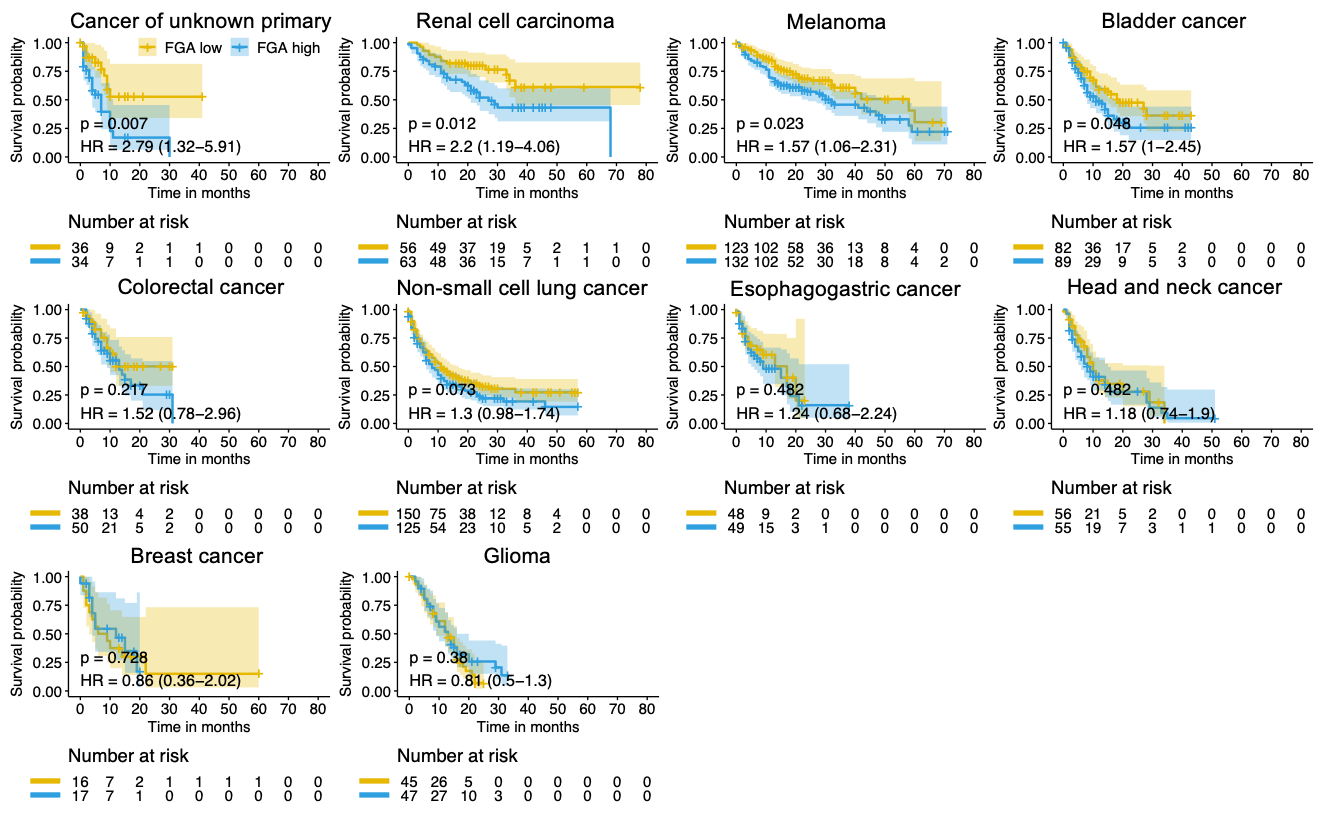

**Supplementary Figure 1. The fraction of copy number alterations (FGA0.2) significantly predicts survival following immunotherapy in four cancer types.** Kaplan-Meier survival curves following immunotherapy are compared for low-TMB patients with high versus low FGA0.2 (binned at 50th percentile) in all 10 individual cancer types. Here FGA0.2 is re-calculated using the new cutoff of |log2 copy ratio| > 0.2. Two-sided log-rank p values are indicated, with univariate Cox regression HR with 95% confidence interval. Data from the Samstein et al. cohort ^1^.

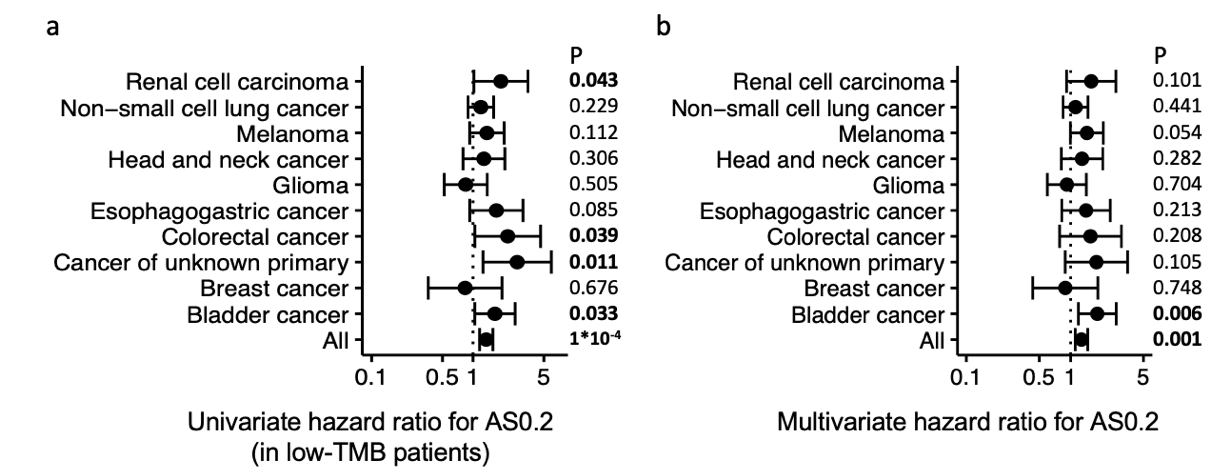

**Supplementary Figure 2. The predictive power of tumor aneuploidy score (AS0.2) in predicting survival following immunotherapy.** (**a)** Hazard ratios (with 95% confidence interval) for AS0.2 in low-TMB patients of different cancer types using univariate Kaplan-Meier survival analysis (AS0.2 binned at the 50th percentile). Two-sided log-rank p values are indicated. (**b)** Hazard ratios (with 95% confidence interval) for AS0.2 of different cancer types using multivariate Cox proportional-hazards regression with AS0.2, TMB, and drug class (AS0.2 binned at the 50th percentile and TMB binned at the 20th percentile). Wald test multivariate adjusted p values are indicated. Here AS0.2 is re-calculated using the new cutoff of |log2 copy ratio| > 0.2. The univariate and multivariate survival analysis are done following ^3^ as is. Data from the Samstein et al. cohort ^1^.

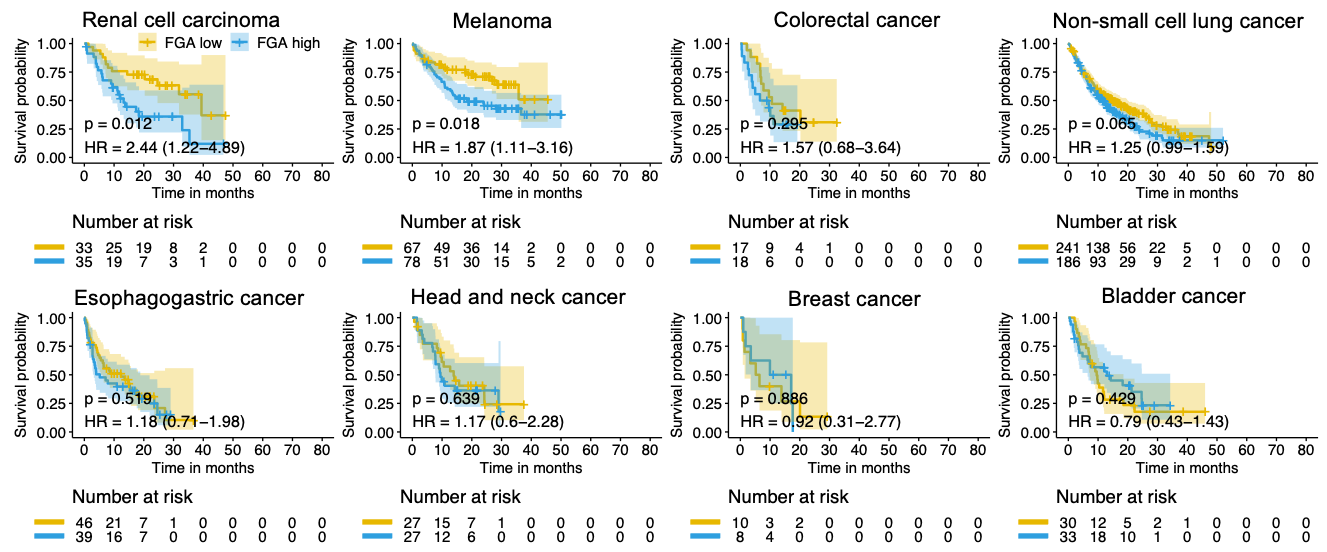

**Supplementary Figure 3. The predictive power of fraction of copy number alterations (FGA0.2) for survival following immunotherapy in another MSK cohort.** Kaplan-Meier survival curves following immunotherapy are compared for low-TMB patients with high versus low FGA0.2 (binned at 50th percentile) in all 10 individual cancer types. Two-sided log-rank p values are indicated, with univariate Cox regression HR with 95% confidence interval. Data from the Chowell et al. cohort ^8^.
